# scSniper: Single-cell Deep Neural Network-based Identification of Prominent Biomarkers

**DOI:** 10.1101/2023.11.22.568389

**Authors:** Mingyang Li, Yanshuo Chen, Jun Ding

## Abstract

Discovering disease biomarkers at the single-cell level is crucial for advancing our understanding of diseases and improving diagnostic accuracy. However, current computational methods often have limitations, such as a reliance on prior knowledge, constraints to unimodal data, and the use of conventional statistical tests for feature selection. To address these issues, we introduce scSniper, a novel approach that employs a specialized deep neural network framework tailored for robust single-cell multiomic biomarker detection. A standout feature of scSniper is the mimetic attention block, enhancing alignment across multi-modal data types. Moreover, scSniper utilizes sensitivity analysis based on a deep neural network for feature selection and uncovers intricate gene regulatory networks without requiring prior knowledge. Comprehensive evaluations on real-world datasets, including COVID-19 CITE-Seq and LUAD scRNA-Seq, demonstrate scSniper’s exceptional ability to identify critical biomarkers consistently outperforming traditional methods like MAST, Wilcox, and DESeq2. The scSniper tool and related experimental codes are publicly accessible at https://github.com/mcgilldinglab/scSniper.

## 1 Introduction

Disease biomarkers serve as invaluable indicators in medical diagnostics, often revealing the presence or progression of pathological conditions [1,2]. The transformative technology of single-cell sequencing has expanded our perspective, allowing researchers to delve into the nuances of individual cell responses and their roles in disease processes [3,4,5,6]. This fine-scale resolution has proven indispensable for biomarker discovery, as it unravels the detailed landscape of cellular heterogeneity, enabling the identification of distinct cellular populations pivotal in disease onset or progression [7]. The pursuit of biomarkers through single-cell approaches thus stands at the forefront of precision medicine, promising a new era of targeted therapies and individualized treatment strategies [8,9].

In recent years, numerous computational tools and methodologies, such as MAST, Wilcoxon, and DESeq2, have emerged, primarily focusing on differential gene expression [10,11,12]. These tools have provided significant insights into cellular behavior and dynamics. However, the inherent complexity of cellular systems means that key genetic elements, particularly oncogenes, often do not simply rely on differential expression. Instead, their influence stems from their ability to modulate other genes within intricate signaling pathways. This underlines the complex web of gene-gene interactions that play a pivotal role in disease onset and progression [13,14,15]. Relying solely on differential expression can thus miss these nuanced interactions, leading to potentially incomplete or even incorrect biomarker identifications. Furthermore, while certain methods have attempted to model such interactions by incorporating prior knowledge [16,17], this approach is not without its limitations. Acquiring accurate and comprehensive prior knowledge, especially for cross-modality interactions, remains a significant challenge in the field [18]. Existing tools, while adept at analyzing individual biological data modality, often fall short in multi-omic biomarker discovery, neglecting the intricate interplay between genomics, proteomics, and metabolomics [19]. This oversight may lead to missed critical biomarkers pivotal for disease understanding. A pressing need thus remains for methods that can holistically integrate and interpret multi-omic data.

Addressing these multifaceted challenges necessitates a fresh approach. Here we introduce scSniper. Unlike many existing methods, scSniper does not base its biomarker identification on differential gene expression. Instead, it harnesses the power of deep neural network sensitivity analysis for feature selection, guaranteeing the identification of the most pivotal gene biomarkers with enhanced accuracy. This technique also presents a groundbreaking mechanism to decipher and capitalize on feature-feature regulatory interactions (i.e., gene interactions), minimizing the dependency on pre-existing knowledge of the gene regulatory network. Furthermore, scSniper’s trailblazing mimetic attention mechanism allows for the fluid integration of varied omics data, ensuring the capture of effective biomarkers across diverse modalities. As a result, its biomarker identification process is both exhaustive and razor-sharp. Rooted in a distinct deep neural network feature selection formulation, our method synergizes feature rankings from multiple modalities, presenting a comprehensive approach to feature selection from all pertinent data sources. Central to scSniper’s novelty in multi-omic biomarker discovery is its utilization of a mimetic attention mechanism within its deep neural network framework. This attention-driven approach enables scSniper to focus on and prioritize key multiomic features across diverse data modalities. Integrating a disease classifier and an autoencoder, scSniper adopts a synergistic method in biomarker discovery, creating disease-focused joint cell embeddings that represent multi-omic cells. This effective multimodal data integration for joint cell embeddings enhances cell clustering and identification accuracy, essential for downstream single-cell level biomarker discovery.

Rigorous evaluations on real-world datasets, including COVID-19 CITE-Seq and LUAD scRNA-Seq datasets, underscore scSniper’s prowess. The results clearly indicate that scSniper outperforms traditional differential expression-centric tools in various metrics, including pathway enrichment, classification, and survival analysis. Notably, in its application to LUAD datasets, scSniper adeptly identifies cancer marker genes, even those with subtle expression variations. Its performance remains consistent across diverse datasets, solidifying its position as a premier tool in biomarker detection.

## 2 Methods

Problem setting Given a multi-omics dataset, encompassing *N* cells, *K* modalities, and *M* features distributed across all modalities, we represent the expression level of feature *m*_*i*_ for each cell within an expression matrix **X** ∈ ℝ ^*N ×M*^ . We also have a patient group label *Y* ∈*G*^*N*^, where *G* denotes the universe of all group labels. Our main goal is to derive a ranking for every feature *m*_*i*_ utilizing a ranking scoring mechanism denoted by *T* = { (*m*_*i*_, *s*_*i*_) | *i* = 1, …, *M*} . We define the score *s*_*i*_ is calculated by adding two components. The first component assesses the importance of a feature by evaluating how changes in its expression can affect the probability of an outcome. The second component focuses on the relevance between different features. Intriguingly, the formulation in equation provides insights into potential model architectures: the first part lays out the foundation of a classifier (Equation 1) dictated by *θ*, whereas the second part elucidates an autoencoder’s structure (Equation 2), governed by *ϕ*. Aligned with this architecture, our scoring mechanism’s design prescribes a definitive training objective, *L*_*i*_, showcased in figure 1(*a*). This objective aims to optimize the likelihood as articulated in equation 3. During the evaluation process, we compute *s*_*i*_ by assigning *m*_*i*_ = 0 and reassess the model via sensitivity analysis as detailed in finding effective biomarkers section.

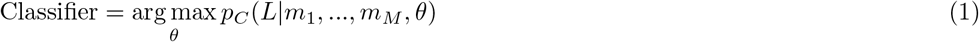

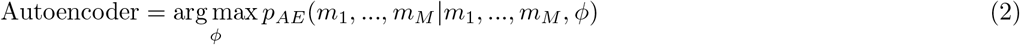

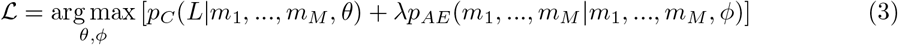

scSniper design Given the expression matrix **X**, we can segregate it into modality-specific sub-matrices 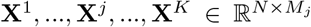, with *M*_*j*_ denoting the feature count for modality *j*. scSniper integrates *K* specialized encoders, *E*_1_, …, *E*_*K*_, to create modality-specific embeddings, termed as **Z**^1^, …, **Z**^*K*^. A custombuilt mimetic attention block aligns these to form a unified embedding, **Z**_*J*_ . This joint representation helps a classifier in pinpointing the originating category of the cell. Concurrently, *k* decoders reconstructs the initial modality input, leading to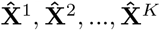 .

**Fig. 1.**
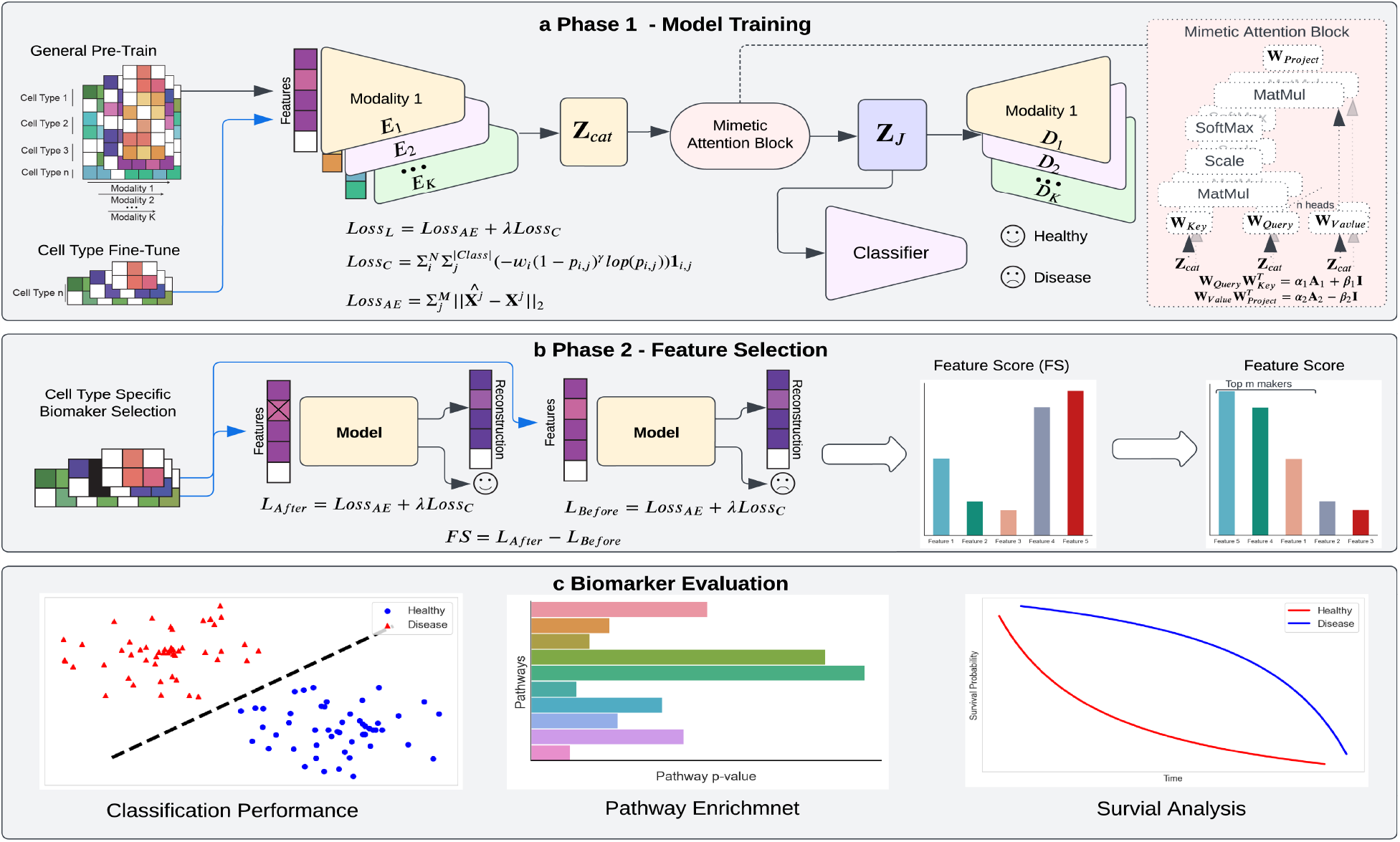
Overview of the scSniper pipeline. (a) This panel outlines our four-stage training strategy: In Stage 0, the autoencoder is warmed-up using the complete dataset. In Stage 1, the autoencoder undergoes fine-tuning specific to a single cell type. During Stage 2, a classifier is trained using data from one cell type while the autoencoder remains frozen. Finally, in Stage 3, both the classifier and autoencoder are trained concurrently. (b) This panel showcases our sensitivity-driven feature selection approach, where features are ranked based on the decrease in accuracy upon their removal. The resulting list of ranked genes serves as potential biomarkers for the specific cell type. (c) This panel highlights the three evaluation strategies we utilized to validate the identified biomarkers

For training optimization, **phase 1**, as shown in equation (3) and figure 1(a), we adopt a staged training approach. In stage 0, the scSniper’s autoencoder is pretrained with the entire dataset, in line with equation (2), to achieve a comprehensive data understanding. It is then fine-tuned with cell type specific data by using inputs of a singular cell type denoted by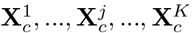, 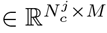, where *c* is the specific cell type and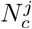represents its cell count. This stage focuses on maximizing the objective as per equation (2). Stage 3 (figure 1(a)) introduces a lightweight classifier module trained from scratch, purposed for classifying the right patient category, optimizing the probability as per equation (1) through the cell type-specific joint embedding **Z**_*c*_. In the stage 4, the culmination of our training strategy involves a combined training of both the autoencoder and classifier, aligning with equation (3). The stage 2 to 4 are repeated for every cell type. For feature selection (figure 1(b)), **phase 2**, we measure the changes in loss by removing specific features and compared with the original value, essentially operationalizing equation (3) via the selectivity analysis technique. Features are subsequently ranked by the degree of loss change, selecting those exhibiting significant fluctuations as potential biomarkers. The network design details are in the supplementary method section.

### Mimetic attention block

Upon using the modality-specific encoders, the model obtains individual modality embeddings. However, due to cellular heterogeneity, these embeddings might remain misaligned. To align these embeddings and find the inter-modality correlations, we incorporate the mimetic attention mechanism. Renowned for its efficacy, especially in scenarios with limited datasets, the mimetic attention [20] consistently surpasses the conventional attention mechanism. This superiority is from its initialization strategy that mirrors patterns observed in extensive dataset training. Before applying the attention, in scSniper, we concatenate **Z**^1^, …, **Z**^*j*^, …, **Z**^*K*^ into **Z**_*cat*_ ∈ ℝ^*N ×d*^, where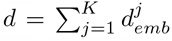 and 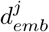 is the embedding dimension for modality *j*. The same is for the cell type specific embedding. Diving into its technicalities, the multihead attention mechanism, illustrated in equation (4), is equipped with the query and key weight matrices, symbolized as 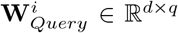 and 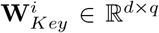 where *q* represents the dimension of each head 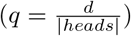and *i* is the head index. The value weight matrix and the projection matrix are **W**_*Value*_ ∈ ℝ^*d×d*^ and **W**_Project_ ∈ ℝ^*d×d*^ respectively.

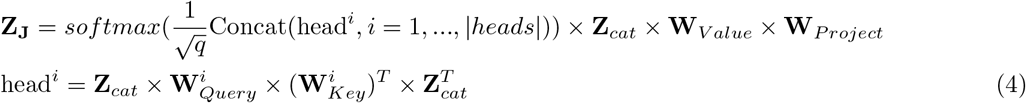

We initialize every query and key matrix the same way. We ignore the superscript *i* from now on. We opt to initialize the weight matrice using equation (5), where each elements of **A**_1_ and **A**_2_ is drawn from (0, 1) and both *α*_1_, *β*_1_ and *α*_2_, *β*_2_ range between [0,1].

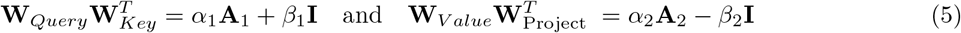

To initialize the individual weight matrix, we applied Singular Value Decomposition (SVD) to equation (5).

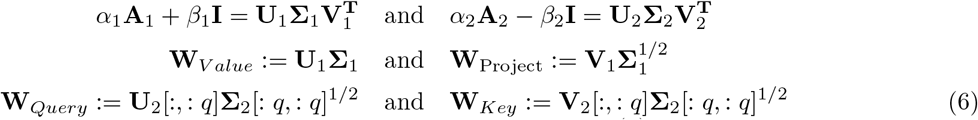

### Optimizing the autoencdoer

Maximizing the likelihood of equation (2) is equivalently to minimizing the negative log likelihood (NLL). We use mean squared error (MSE, equation 7) to approximate the NLL, where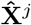 indicates the reconstructed input for modality *j* and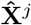 indicates the original input for modality *j*. For cell type specific training, one should replace **X**^*j*^ with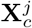, and the same for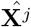

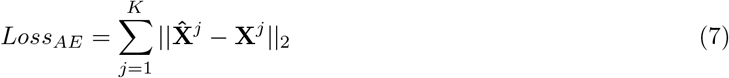

### Optimizing the classifier

Due to the potential class imbalance in between patient categories, we apply focal loss (equation (8)) as the NLL in the classifier m odule, w here *w* _*j*_ is the weight of each classes, *γ* is to focus on hard-to-classify cell, *p*_*i,j*_ is the probability of the class *c* for cell *i*, and (1 − *p*_*i,j*_ )^*γ*^ ensures wellclassified e xamples g ets l ow loss. T he 𝟙 *i,j* is an indicator function and returns 1 w hen t he c lass l abel j is the correct classification for c ell *i* . To reduce the class imbalance, the focal loss shifts the model focus from the well-classified cells to misclassified cells. In the training of the classifier, we firstly keep the autoencoder module weights *θ*_*AE*_ frozen and maximize the likelihood of equation 1. In this way, the classifier is able to learn the probability given the joint embedding. The second stage of the training, we trained both the classifier module *θ*_*C*_ and the autoencoder module *θ*_*AE*_ to further maximize the likelihood of equation 3.

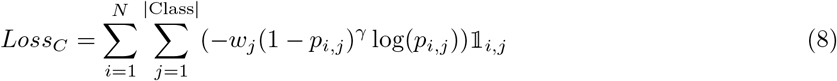

### Finding effective biomarkers

With the autoencoder and classifier defined as per equations (1) and (2), we can reinterpret the term *score*_*i*_ in the context of losses, as delineated in equation (9). The terms *Loss*_*C*_(*m*_1_, …, *m*_*i*_ = 0, *m*_*i*+1_, …, *m*_*M*_ ) and *Loss*_*AE*_(*m*_1_, …, *m*_*i*_ = 0, *m*_*i*+1_, …, *m*_*M*_ ) are deduced by zeroing out feature *m*_*i*_ while preserving the integrity of the remaining features. Iteratively, we evaluate each feature *m*_*i*_, *i* = 1, …, *M* to determine its respective score *s*_*i*_. Subsequently, a descending order ranking is applied, with the top *N*_*F*_ features being designated as the biomarkers, where *N*_*F*_ is a user-specified parameter for the maximum number of selected biomarkers.

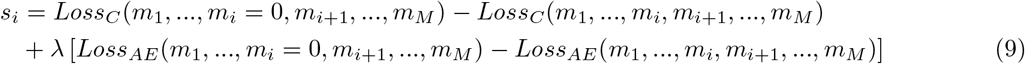

## 3 Experimental Setup

In the realm of single-cell multi-omic biomarker discovery, available methodologies are limited. In this study, we conducted a comparative analysis of the scSniper method against three leading biomarker discovery techniques tailored for single-cell data: Wilcox, MAST, and DESeq2. To ensure a fair comparison, we focused our assessment specifically on the RNA modality since the other benchmarked methods are unimodal. However, it’s essential to note that in multi-omic application scenarios, our method still utilizes the multi-omic input to train the model and identify biomarkers across different modalities. Nonetheless, for the sake of fairness, only the identified gene biomarkers within the RNA modality were subject to comparison with other methods. Each approach was tasked with identifying the top *N*_*F*_ genes for each cell type, with *N*_*F*_ defaulting to 100. Following necessary preprocessing steps, we partitioned all datasets using an 8:2 split for out-of-sample testing and subsequent evaluation.

We benchmarked our methods against other state-of-the-art methods in 4 real and simulated datasets. **Coronavirus Disease 19 Dataset 1 (COVID 1) [21]**: We utilized the **multi-omic** CITE-seq dataset termed COVID 1, which includes 38,652 cells from 18 COVID-19 patients across 24 cell types. These patients were grouped into three categories: ‘Treated Progressive’, ‘Treated Stable’, and ‘Non-treated Stable’. Here, ‘Progressive’ denotes marked COVID-19 progression, whereas ‘Stable’ indicates a recovery phase. Following the methods laid out in the referenced study, we conducted preprocessing. Cells from ‘Progressive’ patients received a ‘disease’ label, while those from ‘Stable’ conditions were designated as ‘healthy’. **Coronavirus Disease 19 Dataset 2 (COVID 2) [22]**: We examined another **multi-omic** CITE-seq dataset dubbed COVID 2, which comprises 647,366 cells across 18 cell types, collected from three distinct COVID-19 testing facilities presenting diverse disease states. From this collection, we focused on 130,644 cells specifically derived from the ‘Cambridge’ center. In preprocessing both RNA-seq and protein (ADT)-seq modalities, we followed the procedures detailed in the original study, but included an additional step to exclude mitochondrial genes. For simplification, we grouped original patient labels like ‘Asymptomatic’, ‘Mild’, ‘Moderate’, ‘Severe’, and ‘Critical’ under a unified ‘disease’ label, while maintaining ‘Healthy’ as a separate category. **Lung Adenocarcinoma Dataset (LUAD) [23]**: We utilized the **unimodal** LUAD dataset, which comprises 208,506 cells from 44 patients spanning stages ‘I’ to ‘IV’ and covers both cancerous and non-cancerous tissues across 50 cell types. The data, primarily RNA modality, underwent preprocessing per the methods delineated in the cited study. Specifically, we removed ribosomal and mitochondrial genes, filtered out cells with fewer than 3 genes or genes present in under 200 cells, normalized using a 1*e*^4^ factor, applied a *log*1*p* transformation, and 178,106 cells remained, with those from cancerous tissues labeled ‘disease’ and those from non-cancerous tissues marked ‘healthy’. **Simulation Dataset**: We utilized a simulated dataset derived from the DOGMA-seq platform [24] to assess biomarker identification against known ground truth genes. Following the preprocessing guidelines established in the source paper, we employed Uniform Manifold Approximation and Projection (UMAP) for cell clustering and subsequently selected the most substantial cluster for our simulation endeavors. Given the RNA-seq modality preference of benchmarking techniques, our investigation was limited to RNA-seq data. Applying filters in line with the recommendations of [25], we concentrated on a subset of 10,000 highly variable genes. Out of this set, 100 genes showing the most non-zero values were designated as biomarker contenders. These selected genes were then split into two distinct groups, and their raw count matrices were subjected to perturbations of incremental multipliers (1.01x, 1.05x, 1.1x, 1.15x, 1.2x, and 1.3x) to emulate subtle variations in gene expression.

Subsequently, we adopted the following strategies to assess the performance of various methods in the identification of disease biomarkers across different cell types: **Pathway Enrichment** In our study, we employed pathway enrichment analysis to validate the biomarkers identified. This is a statistical approach designed to identify pathways that are notably overrepresented in a given gene list, beyond what might occur due to random chance. We utilized the ToppGene Suite [26] for the pathway analysis of our biomarker genes, which shows Reactome, KEGG, and WP pathways. A stronger enrichment of the biomarker list, combined with its biological relevance to the specific disease under study, serves as an indicator of the effectiveness of the identified biomarkers. In our analysis, we relied on the Bonferroni adjusted p-value to account for multiple comparisons. We select top 100 gene markers from each methods. For protein (ADT) markers, we select top 50 protein markers. For cross-modality biomarker discovery, we select top 150 markers. We also applied go term analysis on LUAD dataset. **Logistic Regression** To further validate the efficacy of our predicted biomarkers, we assessed their ability to accurately predict disease outcomes. Specifically, we took the top *N*_*F*_ markers from each method, distinguishing between disease and healthy states, and applied logistic regression on a cell-type-specific basis. This analysis was conducted using the testing data, wherein each cell type was compared against its original class labels. The performance of these selected gene markers was gauged using weighted F1 scores, offering a comprehensive evaluation. These results strengthened our confidence in the selected gene markers’ prowess in distinguishing between various health states across different cell types. **Survival Analysis** For our third validation of the predicted biomarkers, we turned to prognosis analysis, as we believe that an effective set of biomarkers should demonstrate strong prognostic capabilities. The Kaplan-Meier (KM) plot [27] is a widely used survival analysis tool that serves to visualize and assess prognosis outcomes. A signature gene’s expression disparities are reflected on the survival plot, with the significance of these differences quantified by a p-value. In our analysis, we utilized the KM plot to evaluate our biomarker genes specifically for non-small cell lung cancers, employing the tool referenced in [28]. To provide a consolidated assessment of multiple p-values, we calculated the geometric mean of these values. The scSniper multi-omics details are in the supplementary experiment setup section.

## 4 Results

### Uncovering robust multi-omic COVID-19 disease biomarkers with scSniper

We applied scSniper to a multi-omic COVID-19 dataset, termed COVID1. scSniper is intrinsically multi-modal and can handle inputs from various modalities, capitalizing on cross-modal information fusion and interaction. However, in our analysis, we maintained a specific focus. To ensure a fair comparison, given that all other benchmarked methods are unimodal, we also provided the gene biomarkers derived from only the RNA modality by scSniper for comparison (term as scSniper). In Figure 2(a), we presented 20 top-ranked marker genes identified by scSniper, originating from classical monocytes. Notably, the figure showcases an increase in ribosome related genes in expression from classical monocytes derived from diseased patients compared to the healthy control. The importance of RPS14 as a transcription factor for COVID-19 has been supported by its listing as a predicted marker in [29]. Other genes like RPS19, RPL26, and RPS7 have also been flagged as differentially expressed genes in COVID-19 by [30]. Owing to the invasion mechanisms of COVID-19 which leverage a variety of receptors and exploit the host cell’s machinery [31,32,33,34], it is plausible to witness a similarity of biomarkers across various cell types. Expanding upon our exploration of scSniper’s efficacy in RNA biomarker identification, it’s important to highlight the tool’s exceptional multi-omic capability in pinpointing protein biomarkers, supported by concrete examples from our analysis. Within a comprehensive panel of 198 proteins, scSniper adeptly pinpointed top 10 vital protein biomarkers originating from classical monocytes (an example cell type). These protein markers, such as CD32, known for its role in antibody-dependent enhancement through FcRs binding, and CD29, a supporter of SARS-CoV-2 entry, provide invaluable insights into the intricate COVID-19 landscape [35,36]. Additionally, CD224 acts as an immune checkpoint inhibitor, influencing the immune response in COVID-19, while CD19’s expression level is intricately linked to the acquisition of anti-SARS-CoV-2 IgG [37,38]. Furthermore, CD82 has emerged as a reliable COVID-19 biomarker, and CD88 plays a pivotal role in recruiting neutrophils and monocytes in the context of COVID-19 [39,40]. These findings not only reaffirm scSniper’s versatility and effectiveness but also showcase its ability to offer complementary insights into the complex biology of COVID-19, bridging the gap between RNA and protein modalities. A byproduct of scSniper’s design is its delivery of joint cell embeddings, as visualized in Figure 2(b). These embeddings allow for meaningful cell segregation based on their types. For instance, the proximity between classical monocytes and other monocytes (NC and IM) suggests their high similarity, while classical monocytes exhibit a pronounced separation from the B cells.

**Fig. 2.**
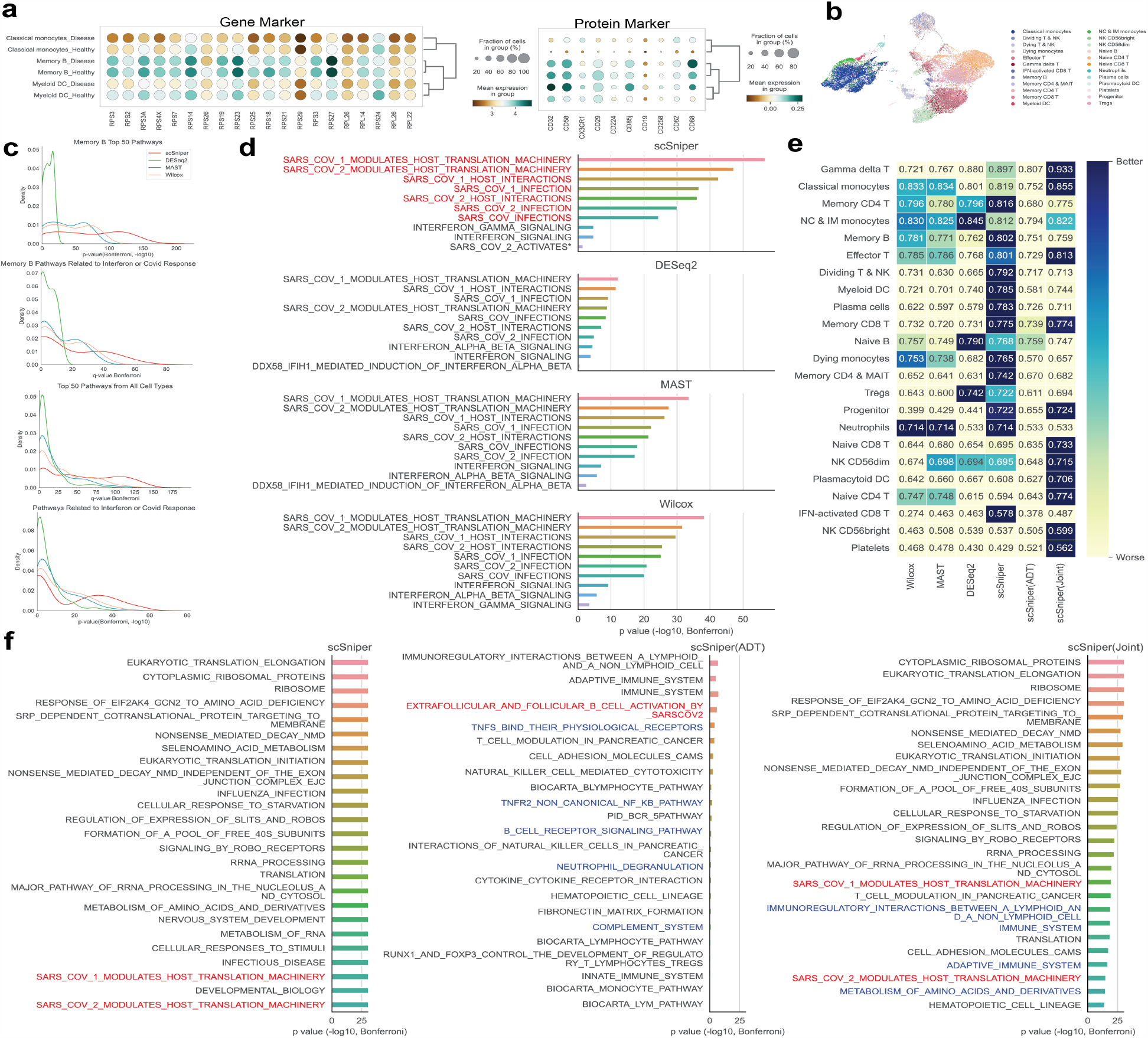
Detailed analysis of the COVID-1 dataset utilizing scSniper. (a) Dot plot comparison of multiomic biomarkers across three immune cell types, contrasting healthy and diseased states. (b) UMAP visualization representing the joint embedding enhanced by mimetic attention. (c) Kernel density estimation (KDE) plots for p-values associated with the top 50 enriched pathways within identified biomarkers. The highest KDE plot illustrates the top 50 pathways in Memory B cells (as an example cell type), followed by a plot detailing COVID-specific pathways in Memory B cells. The third plot encompasses the top 50 pathways across all cell types, and the final plot highlights COVID-specific pathways across all cell types. (d) A bar plot depicting the significance of the top 10 pathways in an exemplary cell type, Memory B cells, as identified by each method. **scSniper(Joint)** represents the integration of multi-omic biomarkers, indicating the joint discovery of gene and protein markers. (e) F1 scores from logistic regression analysis per cell type using biomarkers identified by each benchmarked method. (f) Bar plot detailing the top 25 pathways identified by gene markers, protein markers, and their combination in Memory B cells, as revealed by scSniper. Please note that ‘scSniper” without specification refers to scSniper(RNA).

To delve into the biological significance of gene biomarkers identified by scSniper in the COVID1 dataset, we examined Kernel Density Estimation (KDE) plots that illustrate pathway enrichment under four distinct conditions (Figure 2(c)). The x-axis represents the negative logarithm of the p-value (−log_10_ p-value). The initial plot highlights the top 50 pathways enriched within B cells, a cell type selected for its crucial role in antibody secretion and immune response against viral adhesion [41,42]. *scSniper* identifies marked peak activities, particularly with a significant concentration around 10^*−*125^, demonstrating its ability to capture biologically relevant pathways, in contrast to the peaks observed with Wilcox and MAST at 10^*−*60^ and 10^*−*75^, respectively, and DESeq2, which does not exhibit similar prominence in low p-value regions. Focusing specifically on COVID-19 related pathways, *scSniper* achieves standout peaks at 10^*−*50^, while Wilcox and MAST show peaks at substantially higher p-values of 10^*−*30^ and 10^*−*25^, respectively, and DESeq2 trails behind these figures. A consistent pattern emerges when analyzing the pathway enrichment across all cell types. Figure 2(d) demonstrates that enriched pathways relevant to SARS-CoV and Interferon responses, identified by the biomarkers detected by scSniper, were exceptionally significant. The pathway termed ‘‘SA RS COV 1 MODULATES HOST TRANSLATION MACHINERY’’, while being the highest-ranked by all methods, was identified by scSniper at a p-value of 10^*−*58^, a value significantly lower than those detected by Wilcox, MAST, and DESeq2. Similarly, scSniper’s identification of the ‘‘SARS COV 2 MODULATES HO ST TRANSLATION MACHINERY’’ pathway at a p-value of 10^*−*47^ further highlights its robust detection capability, underscoring its potential to unravel the intricate biological networks affected by COVID-19.

The multi-omic markers detected by scSniper were also validated using external analytical techniques, such as logistic regression. Figure 2(e) presents the F1 scores obtained by employing gene biomarkers from each method to classify the original patient groups. Notably, scSniper (joint), which conducts joint ranking of RNA and ADT markers, stands out by achieving the highest F1 scores for 10 out of the 23 analyzed cell types. This underscores the robust predictive power of the multi-omic biomarkers identified by scSniper. Meanwhile, scSniper (RNA only), which focuses on discovering RNA markers, ranks second by topping the F1 score charts for 9 out of 23 cell types. This performance is still superior to the other benchmarked methods, with DESeq2 claiming the third spot by leading in only 3 cell types. To scrutinize the multi-modality marker detection capabilities of scSniper further, we conducted an in-depth analysis across three distinct cases, as depicted in Figure 2(f). Initially, we assessed the quality of gene markers on B cells to establish a baseline, as explicated in prior sections. A comparative analysis was performed on the top 25 pathways identified in each case. In the second scenario, highlighted in the middle graph, the pathways enriched with ADT markers were investigated. Notably, the pathway “EXTRAFOLLICULAR AND FOLLICULAR B CELL ACTIVATION BY SARS-COV-2” was identified at the fourth position, unequivocally confirming the presence of COVID-19. The ADT modality unveiled additional COVID-19 related pathways that remained undetected in RNA modality. For instance, several TNF-related pathways were discerned, including “TNFS BIND THEIR PHYSIOLOGICAL RECEPTORS” and “TNFR2 NON-CANONICAL NF-KB PATHWAY.”

[43,44] Furthermore, “B CELL RECEPTOR SIGNALING PATHWAY” corroborated the viral invasion via receptor binding [45,46]. Pathways like “NEUTROPHIL DEGRANULATION” [47] and “COMPLEMENT SYSTEM” [48] were also related to COVID-19. These findings underscore scSniper’s efficacy in discerning biologically meaningful pathways in ADT modality. In the third scenario, scSniper demonstrated its proficiency in jointly identifying multi-modality biomarkers, where it didn’t opt for top RNA and ADT markers separately, but instead ranked them jointly by their scores. The resulting pathways presented a more comprehensive list, with the top 15 pathways being identical to those identified by RNA markers alone. Moreover, the ranking of COVID-related pathways improved, with several of them ascending in the list. New COVID-related pathways, such as “IMMUNOREGULATORY INTERACTIONS” [49], “IMMUNE SYSTEMS” [50], and “METABOLISM OF AMINO ACIDS AND DERIVATIVES” [51] were also unveiled. This denotes that the integration of jointly discovering RNA and ADT marker enhances the comprehensiveness and relevance of the results. We also benchmarked the methods on another dataset, COVID2. Preliminary results show scSniper’s superiority in all considered metrics. For detailed results, refer to the supplementary Figure S1.

### scSniper effectively identify disease biomarkers in LAUD single-cell RNA-seq data

To showcase the potency of scSniper, especially when juxtaposed against unimodal datasets, which remain quite prevalent, we elected to benchmark our method against a single-cell LUAD cancer RNA-seq dataset. The exceptional prowess of scSniper in discerning biologically pertinent marker genes from the LUAD cancer dataset becomes strikingly evident. This capability transcends mere detection of differential expression, zeroing in on genes bearing profound functional implications. One of scSniper’s key strengths lies in its refined sensitivity, adeptly picking out genes even with nuanced expression differences but having significant biological ramifications. A meticulous dot plot, as seen in Figure 3(a), elaborates on the expression behaviors of the top 20 lung cancer-related marker genes within tumor endothelial cells (ECs), as identified by scSniper. Distinct markers reveal varying expression dynamics across tumor ECs, tip-like ECs, and stalk-like ECs. We choose ECs as examples as it plays vital roles in lung cancer initialization, progress, and metastasis [52,53]. Noteworthy is the SPARC gene, characterized by its muted expression in tumors, yet playing a pivotal role in microvascular remodeling [54,55]. On the other hand, the CAV1 gene, notorious for bolstering lung cancer cell proliferation, manifests heightened expression during pathological conditions [56]. Other markers, such as HSPB1, EGFL7, and PPIA, while not overtly differentially expressed, resonate strongly with their established biological significance [57,58,59]. Figure 3(b) serves as an emblem of scSniper’s autoencoder module’s finesse in sculpting biologically coherent cell embeddings, facilitating a precise delineation of diverse cell type clusters.

**Fig. 3.**
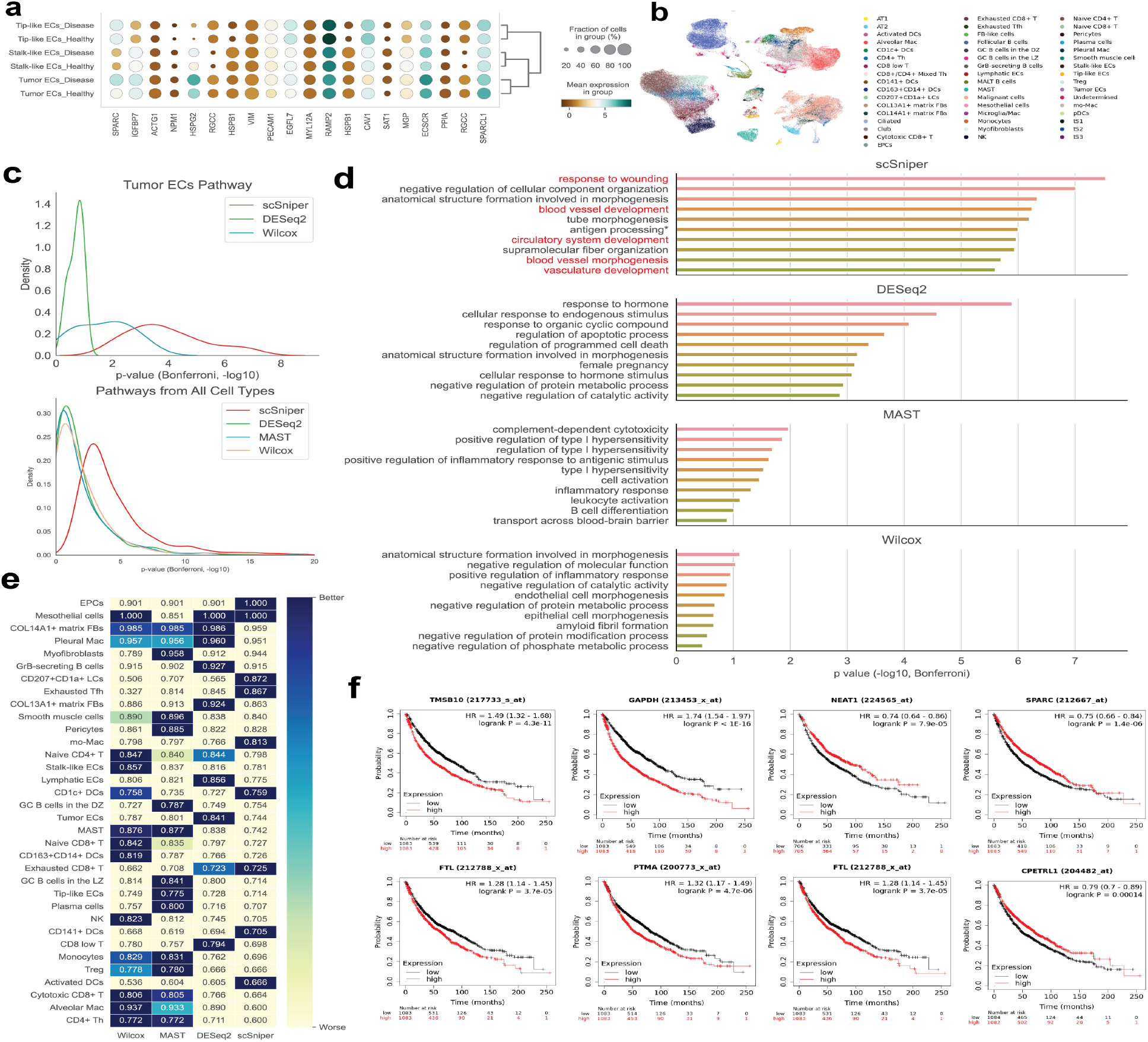
Analysis of LUAD dataset showcasing the efficacy of the scSniper method. (a) Dot plot comparison of three ECs in both healthy and disease states. (b) UMAP visualization reflecting the joint embedding post mimetic attention processing. (c) Dual KDE plots depicting p-values of the top 50 pathways for cell types, with the x-axis scaled to *−* log_10_ p-value. The upper Kernel Density Estimation plot represents top 50 pathways from Tumor ECs, while the lower one depicts pathways across all cell types. (d) Bar chart representation of p-values for the leading 10 biological processes pinpointed by various methods, specifically for tumor ECs. (e) F1 score determinations for each cell type using logistic regression across different methods. (f) Kaplan-Meier (KM) survival plots, indicating the capability of the identified biomarkers in prognosis prediction.

From the vantage point of pathway enrichment tied to identified biomarkers, the supremacy of scSniper stands illuminated in the LUAD lung cancer dataset. A KDE in Figure 3(c) delineates the p-values associated with various pathways enriched with the identified biomarker genes. Upon assessing the top 50 pathways across all cell types, the eminent stature of scSniper over other conventional methods becomes unequivocal: its peak in the KDE markedly surpasses that of its counterparts. This stark differentiation accentuates the substantial statistical relevance of pathways linked to the biomarkers discerned by scSniper. Further insights are rendered in Figure 3(d), spotlighting the top 10 GO term biological processes (GOBPs) identified in tumor ECs by each method under scrutiny. For clarity, the x-axis is depicted as ( −log_10_ p-value). Amongst the ensemble, scSniper persistently takes the lead in terms of p-value performance. Explicitly, scSniper registers an unparalleled geometric average p-value of 9.53×10^*−*7^. In juxtaposition, DESeq2, the closest competitor, logs an geometric average p-value of 5.43 ×10^*−*4^, with Wilcox and MAST trailing considerably at 0.18 and 0.048, respectively. Impressively, scSniper’s geometric average p-value outperforms DESeq2’s by three magnitudes, trounces Wilcox’s by six magnitudes, and eclipses MAST’s by five magnitudes. Beyond mere statistical distinctions, scSniper demonstrates a unique acumen in unearthing biological processes intrinsic to lung cancer. A notable case in point is the “response to wounding” process, echoing the parallels between cancer growth and wound healing dynamics [60,61].

In our continuous endeavor to further validate the identified biomarkers on the LUAD dataset, Figure 3(e) depicts the F1 scores garnered by deploying biomarkers from each method for patient classification through logistic regression. Within this context, scSniper attains performance metrics that are on par with other state-of-the-art methods benchmarked. This underscores its efficacy in biomarker discovery and disease prediction, demonstrating its strength even in scenarios involving a single modality.

In our pursuit to further validate the effectiveness of the identified biomarkers, we employed Kaplan-Meier (KM) survival plots to scrutinize the top 20 genes pinpointed by each of the four benchmarked techniques, as shown in Figure 3(f). Our primary focus was on assessing the power of these identified biomarkers in prognostic analysis (denoted in survival log-rank test p-values [62]), for which we derived the geometric mean across the top 20 genes (other cutoffs such as 5 and 10 were also used and return similar results). Remarkably, scSniper outshone its peers across all categories. For instance, within the top 20 genes spectrum, scSniper logged a geometric mean p-value of 0.00011, dwarfing the next best, DeSeq2, which marked at 0.00036. Selective KM survival plots representative of scSniper’s adeptness are portrayed in Figure 3(f). Dominantly, scSniper maintained an exceptionally low geometric average p-value, hovering around 1×10^*−*4^. Notably, genes such as GAPDH posted a staggering p-value of less than 1×10^*−*16^, while TMSB10 stood at 4.3× 10^*−*11^. A significant chunk of these top-tier 20 genes bear direct relevance to lung cancer pathology, such as GAPDH’s link to tumor metastasis [63] and TMSB10’s correlation with enhanced lung cancer progression [64].

### Benchmarking using simulation data with groundtruth

To assess scSniper’s ability to detect biomarkers with subtle gene expression changes in the absence of established ground truth markers for real datasets, we utilized a synthetic single-cell RNA (sc-RNA) dataset. This dataset was constructed with 100 genes identified as biomarkers from a total gene pool of 10,000. These biomarkers were split into two groups: the first group had raw counts scaled by factors from 1.01× to 1.3× as depicted in Figure 4(a), while the counts for the second group were unchanged. Analysis revealed a consistent direct correlation between the extent of expression level variations and the number of genes detected by all methods. Notably, scSniper consistently excelled beyond its peers, regardless of the magnitude of expression changes. For example, with a slight increase of 1.05×in raw counts, scSniper detected 16 out of the 100 biomarker genes, with a notably low p-value of 1.1×10^*−*16^, demonstrating its exceptional sensitivity to minimal expression alterations. In stark contrast, other methods like DESeq2, MAST, and Wilcox only managed to detect 8, 2, and 3 genes, respectively. Further, Figure 4(b) substantiates scSniper’s superior performance by charting the number of implanted marker genes against the levels of expression changes. This curve indicates that scSniper consistently outperformed competing methods, particularly when discerning implanted marker genes during minor perturbations where the signals were faint.

**Fig. 4.**
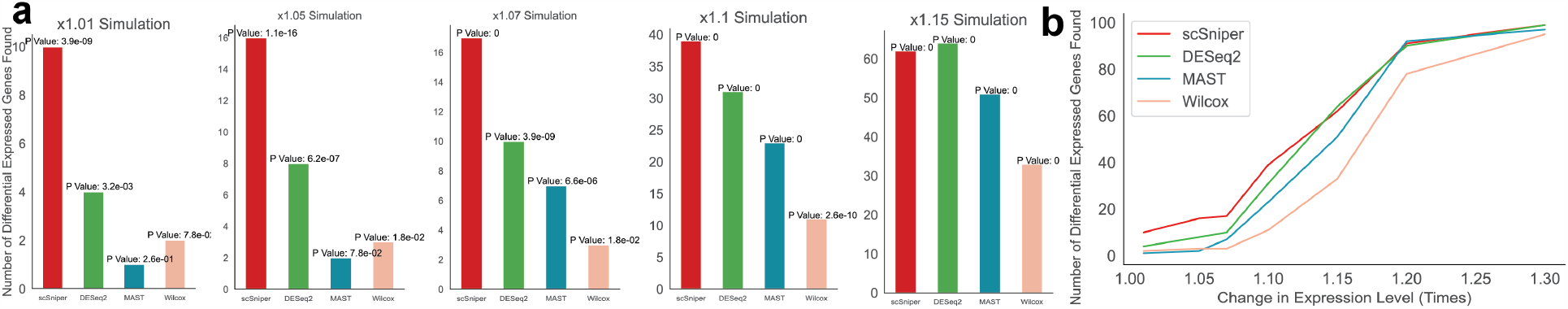
Benchmarking using simulated data with grund-truth (a) Depicts the detection of 100 implanted marker genes within the simulation, quantifying the efficacy of each method, with p-values derived from the hypergeometric test.(c) Provides a comprehensive comparison of biomarkers identified by the four methods across various perturbation intensities employed in distinct simulation scenarios.

## 5 Discussion and conclusion

In the rapidly advancing realm of precision medicine, the identification of disease biomarkers stands as a cornerstone for accurate diagnostics. While single-cell sequencing has revolutionized our understanding of individual cell responses, current computational methodologies, such as MAST, Wilcoxon, and DESeq2, have predominantly focused on differential gene expression, often overlooking the intricate web of gene-gene interactions pivotal to disease onset and progression. More importantly, most existing biomarker detection methods remain unimodal and thus lack the capability to identify comprehensive multi-omic biomarkers across modalities. This lacuna in the existing landscape underscored the need for an innovative approach that goes beyond differential expression and a single modality. Addressing this gap, we introduced scSniper—a novel deep neural network framework tailored for robust single-cell multi-omic biomarker identification. Distinctively, scSniper eschews the conventional differential gene expression paradigm, opting instead for a sensitivity analysis for feature selection, ensuring the pinpointing of critical gene biomarkers with unprecedented precision. Our comprehensive evaluations on real-world datasets have showcased scSniper’s superior performance over traditional tools, emphasizing its effectiveness in biomarker discovery.

The novelty of scSniper stems from several innovative features. Firstly, and perhaps most critically, is its departure from the traditional reliance on differential gene expression. By employing sensitivity analysis, scSniper offers a more refined and precise method for identifying critical gene biomarkers even with subtle expression changes across biological conditions. Secondly, scSniper stands out for its capability in multi-omic biomarker discovery, not just limiting itself to unimodal data. This multi-omic prowess is further enhanced by the incorporation of the pioneering mimetic attention mechanism, adeptly integrating different omics data and offering a more comprehensive biomarker landscape. Lastly, the method’s independence from prior knowledge of the gene regulatory network is a significant advantage. By eliminating this dependency, scSniper bypasses the limitations and challenges associated with obtaining accurate and exhaustive prior knowledge, especially pertinent to cross-modality interactions.

The innovative attributes of scSniper hold substantial promise for the scientific community, particularly in the single-cell biomarker discovery sphere, where it stands to revolutionize disease diagnostics. This tool’s capability to pinpoint subtle yet significant multi-omic markers could dramatically enhance the precision of disease diagnostics, granting clinicians a more detailed understanding of disease mechanisms. Such advancements pave the way for tailored therapeutic interventions and a shift toward personalized medicine. As a vanguard in single-cell biomarker identification, scSniper distinguishes itself with a unique approach that has proven robust across diverse datasets, establishing it as an indispensable asset in the pursuit of effective biomarkers and a deeper grasp of disease pathology.

## Software availability

scSniper, tailored for single-cell biomarker identification, is crafted using Python. The tool is publicly accessible on GitHub at https://github.com/mcgilldinglab/scSniper. The repository provides comprehensive instructions, thorough documentation, and illustrative examples for users.

## Acknowledgements

This work was funded in part by grants awarded to [JD]. We gratefully acknowledge the support from the Canadian Institutes of Health Research (CIHR) under Grant Nos. PJT-180505; the Funds de recherche du Qué bec - Santé (FRQS) under Grant Nos. 295298 and 295299; the Natural Sciences and Engineering Research Council of Canada (NSERC) under Grant No. RGPIN2022-04399; and the Meakins-Christie Chair in Respiratory Research.

